# Rational Design of Selective IL-2-based Activators for CAR T Cells Using AlphaFold3 and Physics-Informed Machine Learning

**DOI:** 10.64898/2026.03.10.710391

**Authors:** Zakaria L. Dahmani, Anupam Banerjee

## Abstract

Recombinant human Interleukin-2 (rhIL-2, Aldesleukin) is used in immunotherapy for metastatic melanoma and renal cell carcinoma. Low-dose IL-2 has been investigated for administration after adoptive T cell transfer to enhance CAR T expansion and sustain effector function. However, systemic IL-2 can cause severe toxicities and promote expansion of regulatory T cells (Tregs). Previous attempts at mitigating cytokine-mediated side effects involved isolating CAR T cell signaling from endogenous immune responses by developing IL-2/IL-2Rβ based selective ligand–receptors systems. Expressing these variant orthogonal (ortho)IL2-Rβ receptors in CAR T cells and supplying variant orthoIL-2, was shown to dramatically improve selectivity in CAR T cell expansion and anti-tumoral potency in a leukemia mouse model. This study describes the computational design of synthetic orthogonal cytokine receptor-ligand systems based on the scaffolds of the human canonical IL-2 and IL-2Rβ. Leveraging state-of-the-art AlphaFold3 (AF3) structure prediction capabilities and a physics-informed constrained sequence generator (CSG), the pipeline generates, filters and ranks sets of putative orthoIL-2/orthoIL-2Rβ mutant designs. Variants displaying minimal predicted off-target interactions and enhanced in target contacts are prioritized for structural modelling. Top designs showed outstanding AF3 structural and interfacial quality metrics ipTM and pTM, with averages between cognate pairs of 0.724±0.05 and 0.770±0.042, respectively. All in-silico hits showed ipTM <0.5 for non-cognates, indicating a good likelihood of orthogonality. Additionally, putative hits showed high levels of predicted structural fidelity to wild-type (WT) human IL-2/IL-2Rβ (PDB: 2ERJ), with an average structural root-mean-square deviation (RMSD) of 0.843±0.375 Å. These mutants incorporated 7–26 interfacial mutations derived from multiple interface selection strategies. Altogether, the results support the putative foldability and selective affinity of top-ranking mutants displaying metrics close-to or within experimental reference range. Finally, strengths and limitations are discussed, alongside the experimental implications of coupling a constrained protein design pipeline to the discovery and validation of selective binders based on naturally occurring scaffolds.

## Introduction

Chimeric Antigen Receptor (CAR) T cell therapy is a groundbreaking cancer treatment that engineers T cells to selectively target and eliminate antigen-expressing cells. CARs combine extracellular antigen-recognition domains with intracellular T cell receptor (TCR) signaling modules, integrating with endogenous pathways to drive cytokine production, proliferation, and tumor-cell clearance. While highly effective in B-cell malignancies such as anti-CD19 and anti-BCMA therapies, CAR T cells face significant limitations, including severe toxicities, restricted efficacy beyond B cells, and complex interactions with the tumor microenvironment.^1–5^ With a total of 23 FDA-approved cytokine products from the CDER-approved Biologic Products list, cytokine-based therapeutics are a well-studied and increasingly popular biologic. IL-2 as the only member of common γ-chain cytokine family approved by FDA, has been investigated in combination with CAR-T therapy for intravenous or subcutaneous administration to treat cancers in initial clinical trials (NCT00019136, NCT03098355, NCT00012207), and was found to promote the expansion of adoptive immune cells in vivo^6,7^. However, the immune stimulation and inflammation due to systemic IL-2 supply can trigger broad immune activation leading to side effects including, but not limited to, cytokine release syndrome (CRS) and neurological complications, ranging in severity from mild to life-threatening (**Fig. 1a**). Altogether these opportunities and challenges have motivated efforts to engineer biomolecular systems that elicit precise T cell responses while minimizing systemic toxicity^5,8–10^.

**Figure 1.**
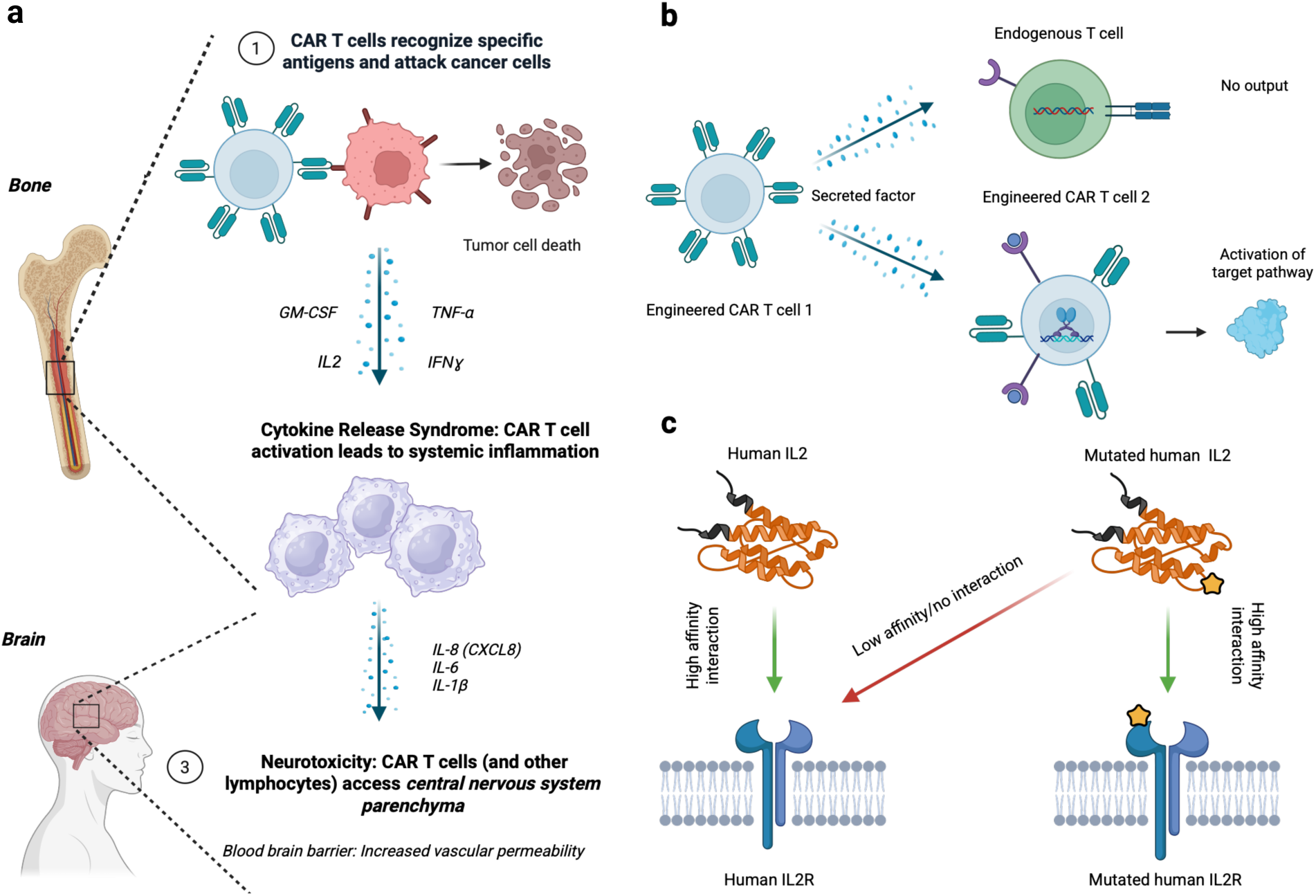
Orthogonal cytokine signaling to control CAR T cell activation and toxicity. **a**, Schematic illustration of cytokine release syndrome (CRS) and neurotoxicity arising from uncontrolled CAR T cell activation and systemic cytokine signaling. **b,** Conceptual representation of orthogonal cytokine receptors and soluble cytokine factors engineered on CAR T cells, enabling pathway activation specifically in modified cells, while endogenous cells lacking the engineered receptor remain unresponsive. **c,** Protein–protein interaction (PPI) view of the orthogonal cytokine system, in which green arrows denote productive interactions between engineered cytokines and receptors, and red arrows denote the absence of interaction with endogenous counterparts.

Orthogonal cytokine–receptor systems provide a promising strategy to enhance CAR T cell safety and specificity (**Fig. 1b-c**). Sockolosky et al.^11^ engineered murine orthogonal IL-2–IL-2Rβ complexes to selectively activate T cells while minimizing off-target effects. Introducing point mutations (H134D, Y135F) in IL-2Rβ and screening IL-2 variants, the authors generated orthoIL-2 variants that expanded orthoIL-2Rβ T cells in vivo in a B16-F10 melanoma model while reducing CD4⁺ regulatory T cells, yielding potent, selective anti-tumor responses. Building on this, Zhang et al.^12^ developed a human-specific ortho-hIL-2/ortho-hIL-2Rβ system that selectively stimulates engineered CAR T cells without activating endogenous populations. In preclinical B-cell leukemia xenograft models, ortho-hIL-2 enhanced CAR T cell engraftment, proliferation, and antitumor efficacy— achieving up to 1000-fold expansion—without systemic toxicity. This approach preserves cytokine functionality at low doses, enables tunable T cell modulation, and may eliminate the need for lymphodepletion, providing a versatile platform for safely controlling T cell activity^12^.

Achieving fully orthogonal cytokine–receptor interactions challenge conventional protein engineering and requires computational approaches capable of estimating sequence– structure–function relationships including binding ability and specificity. Based on classical Maximum Entropy (MaxEnt) argument^13^, the Potts model has been widely and successfully used to design mutant proteins from wild-type (WT) templates and even for early structure prediction attempts^14–19^. Identifying variants that preserve structural integrity while introducing limited sequence variation to narrow experimental sequence space is a well-established strategy to generate focused sets of functionalized analogs based on naturally occurring scaffolds^16,20–22^. Conventional approaches, however, often produce large libraries, yielding few correctly folding proteins and even fewer meeting orthogonality constraints^23^, and rely on computationally intensive procedures, requiring hundreds of Markov Chains (MC) to run in parallel^20^. Recent breakthroughs combining Transformer architectures with Denoising Diffusion Probabilistic Models^24^ (DDPM) have achieved unprecedented structural accuracy by learning to map the distribution of protein sequences to the distribution of structures^25–34^. Trained on the large and trusted structural repertoire of the Protein Data Bank (PDB), these models generate structural hypotheses and metrics characterizing the expected (i.e. predicted) quality of their predictions. Yet even these advanced models, best exemplified by AlphaFold3 (AF3) and RosettaFold Diffusion (RFD), do not support (poly)orthogonality or (poly)affinity constrained design: AF3 is primarily a structure prediction model given some input sequence, and RFD’s conditional design capabilities, though extended, do not include generalized orthogonality^32^.

Developing selective orthogonalized cytokine–cytokine receptor signaling systems represents a critical step toward engineering safer and more controllable orthoIL2 potentiated CAR T cell systems. Conventional cytokine signaling via the *IL-2–IL-2R* and *IL-15–IL-15R* pathways drives potent immune activation but also contributes to severe CRS, neurotoxicity, and recruitment of immunosuppressive regulatory T cells (Tregs). These systemic toxicities, due to uncontrolled engagement of endogenous Jak–Stat cascades regulating multiple immune cell types, impedes the usage of IL2 therapy to potentiate CAR T cell therapy^5,8–10,12,35,36^. On the other hand, orthoIL-2 based systems not only retain the signaling capacity required for CAR T cell persistence and function but significantly increases expansion and anti-tumoral activity while remaining orthogonal to WT IL-2Rβ expressing endogenous cells (**Fig. 1b-c**). The selective activation of CAR T cells with orthoIL2 is a promising strategy for safer and more potent cell-based immunotherapies with reduced systemic impact.

This study describes a computational pipeline geared towards the discovery of focused sets of putative *ortho*IL-2–IL-2Rβ immune signalling pairs. Sockolosky et al.^11^ showed that in order to obtain one viable orthogonal IL-2–IL-2Rβ system by random PCR-based site-directed mutagenesis of the interface region of IL-2 coding sequence, one would need to experimentally generate and explore a space scaling in the ∼10^8^ sequences. By exploring some of the top in-silico hits, emphasis is put on high quality putative *ortho*IL-2–IL-2Rβ AF3 models derived from variant sequence sets of 100-500 candidates. Finally, we discuss how in-silico evaluation and prioritization of mutants could help direct subsequent experimental inquiry.

## Results

### Selective Binder Design with AlphaFold3 and Physics-Informed Machine Learning

To rationally design cytokine systems with minimized off-target activity, we introduce the ICP (Integrative Computational Protein) Design framework, a multi-constraint, data-driven protein design pipeline. Conceived to address current methodological limitations in constrained protein design by extending user control to the protein-protein interaction (PPI) network of designed binders. This computational strategy produces selective binder proteins based on naturally occurring scaffolds, likely to conserve most wild type (WT) properties, while selectively rewiring their PPI network through the introduction of localized mutations (**Fig. 1b-c**). Our framework integrates state-of-the-art AlphaFold3 for the structural prediction of candidate mutants, physics-informed machine-learning for sequence generation, and a rational approach to metric selection and thresholding for the evaluation of constraints satisfaction (**Fig. 2a-b**). Functionally, the pipeline operates as a modular computational funnel: sequences generated by the *Constrained Sequence Generator (CSG)* are filtered through successive tiers of constraint-satisfaction and orthogonality evaluation (**Fig. 2b**), yielding a focused set of high-likelihood design candidates (see **Methods** for implementation details).

**Figure 2.**
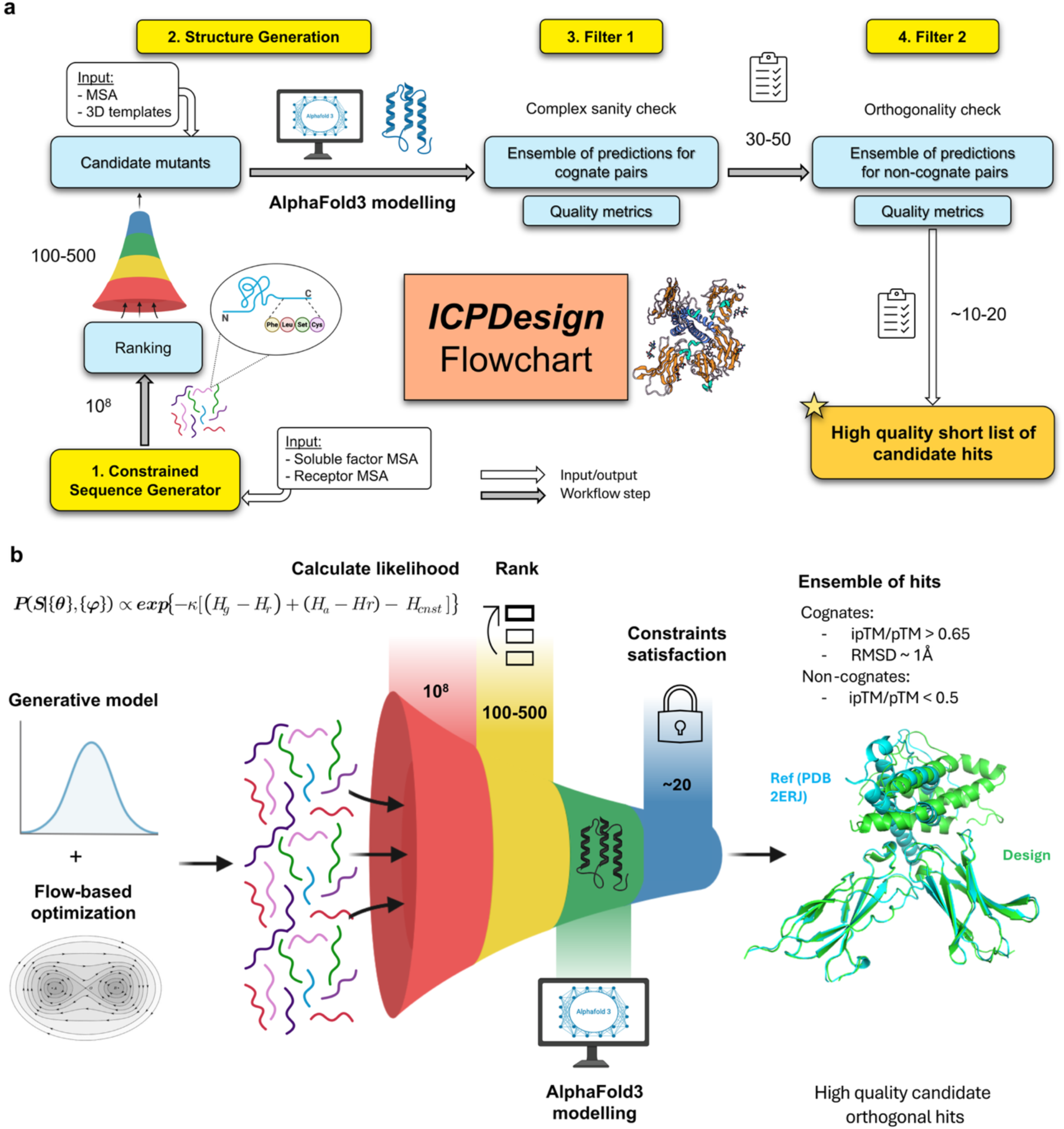
Overview of the ICPDesign framework for selective binder design. **a**, Flowchart representation of the pipeline. Starting from naturally occurring protein scaffolds, sequences are generated by the Constrained Sequence Generator (CSG) and evaluated using a multi-constraint strategy that integrates AlphaFold3-based structural prediction, physics-informed sequence generation and filtering, and rationally chosen metrics with defined thresholds to control protein–protein interaction (PPI). **b,** Funnel representation of the same pipeline, in which large sets of candidate sequences are produced after model training and progressively filtered through successive tiers of constraint satisfaction and orthogonality assessment, yielding a focused set of high-likelihood design candidates.

### Rational Metric Selection and Thresholding to Evaluate Constraints Satisfaction

Following model training and data generation, a large set of proposal mutants is produced using one of or a combination of 3 different modes of sampling (**Methods** and **Table 1**). Log likelihood estimates of proposal mutants are calculated and generated sequences are ranked. Generally, variants in the top 100-500, depending on the model, are picked (**Table 1**). These sequences present outlier values of log likelihood of constraints satisfaction and are expected to yield computational hits (**Methods**). The selected top scoring sequences are then modelled using AF3, with default parameters and independent multiple sequence alignment (MSA) generation for each sequence. AF3 produces ensembles of structural models for which global and local quality metrics will be computed. Using AF3 metrics, predicted structures undergo a third step, where hard thresholds are imposed.

**Table 1.**
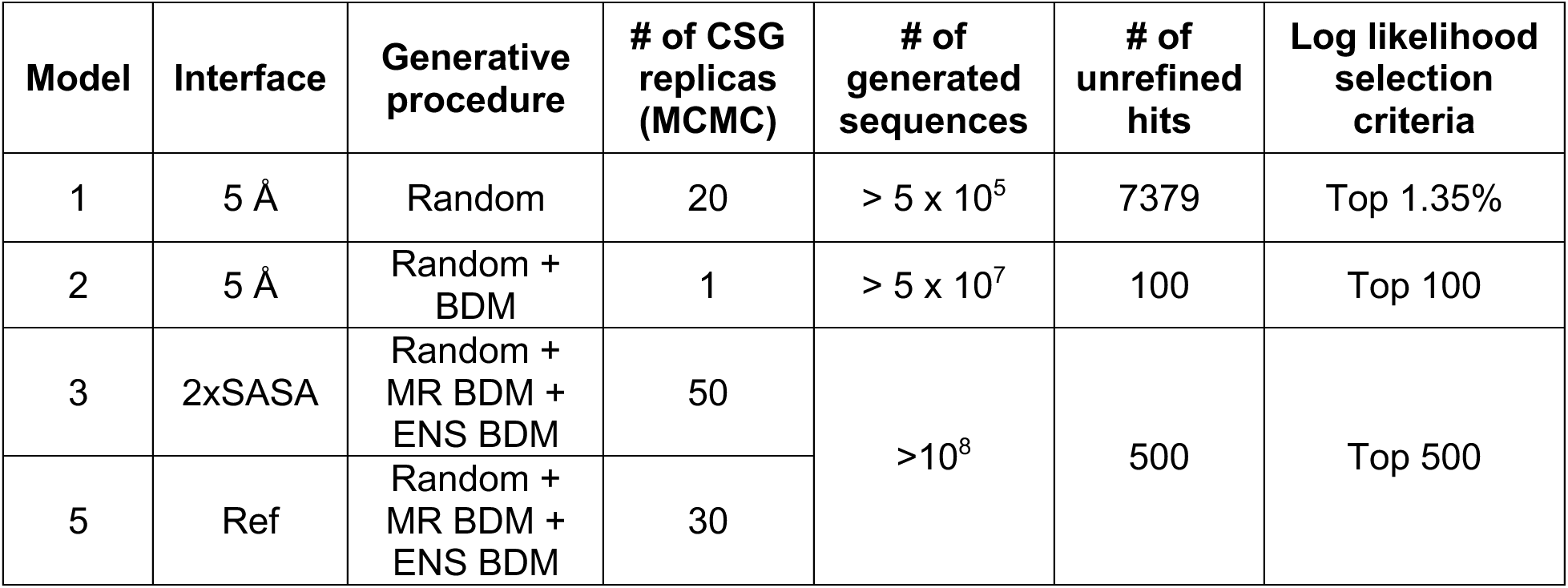
Summary of the generative strategies used to produce orthoIL-2–IL-2Rβ systems using the ICPDesign framework. BDM: Boltzmann-Dirichlet-Multinomial distribution. MR: Multi-replica. ENS: Ensemble. Ref: Reference interface.

**Table 2.** Summary of the interfaces investigated for the computational designs of orthogonal IL-2–IL-2Rβ systems.

| Interface | Retrieval method | WT composition | Number of residues |
| --- | --- | --- | --- |
| 5 Å | Residues whose heavy atoms are within 5 Å of binder. | **IL-2RB:** R42, R41, S69, Q70, K71, L72, T73, T74, V75, H133, Y134, E136, H138, Q188.<br>**IL-2:** L12, Q13, E15, H16, L19, D20, M23, D84, S87, N88, V91, I92, E95 | 27 |
| 2xSASA | $\Delta rSASA \geq 15\%$ + $\Delta SASA > 1\text{ Å}$ upon complex formation | **IL-2RB:** R41, R42, Q70, T73, T74, V75, H133, Y134, Q188<br>**IL-2:** L12, E15, H16, L19, D20, M23, D84, N88, V91, E95 | 19 |
| Ref | Described by Sockolosky et al.<sup>11</sup> | **IL-2:** E15S, H16Q, L19V, D20L, Q22K, M23A, R81D, R83<br>**IL-2RB:** H133D, Y134F. | 10 |

Mutants whose predicted structures show pTM >0.65 and ipTM >0.65, are retained (**Fig. 2b**), The predicted Template Modelling score (pTM), is a quantitative criterion for protein topology classification, i.e. protein pairs with a TM-score >0.5 are mostly in the same fold^37^. In the AlphaFold3 framework this metric estimates the expected TM score of the model to the hypothetical true structure^25^. In addition to assessing the ability of the complex to fold, the ipTM quality metric is used as a proxy to the binding affinity between the two proteins^25,38^. Similar thresholds have been reported in a recent AF3 experimental dataset benchmarking, where pTM/ipTM scores > 0.7 were considered good predictors of structural accuracy^39^. Orthogonality of unrefined hits from the previous step to WT hIL-2Rβ is estimated in the fourth step. Using AF3, the structure of the complex formed by the mutant candidate and the WT hIL-2Rβ is predicted. Abrahamson et al. showed that ipTM scores (interface predicted template modelling, a metric estimating the quality of the predicted interface region between interactor) do correlate with the DockQ score, a *state-of-the-art* structural quality metric of docking models for protein binding. ipTM measures will be used as a proxy to the ability of the mutants to bind non-cognates^25,38–40^. A hard threshold of ipTM<0.5 will be imposed to select the proposal orthogonal mutants (**Fig. 2b**).

To further assess the quality and relevance of the designed IL-2–IL-2Rβ interface variants, we established a set of comparative standards and implemented additional structural and sequence-based constraints within the design workflow. These measures ensured that observed improvements in binding score proxies and interface stability (i.e. ipTM) were meaningful relative to biologically grounded baselines. We defined two main comparative standards to contextualize design performance, and two secondary prioritization constraints: **1. Native reference complex**: the experimentally determined human (h)IL-2–IL-2Rβ structure^41^ (PDB: 2ERJ) served as the baseline model (**Fig. 3a**). We computed C-alpha (C𝛼) RMSDs of high-quality designs with respect to this reference complex and ruled that successful designs should show RMSD ∼ 1 Å. The goal of this soft threshold is to ensure that designs with favorable pTM/ipTM values do not deviate too much from the WT conformation with the aim of increasing the chances of maintaining the template’s original biological function. An AF3 model of the (h)IL-2–IL-2Rβ complex was predicted and its pTM/ipTM values were reported and compared to pTM/ipTM values of proposed designs (**Fig. 3b-c**), ideal candidate hits would show pTM/ipTM values close to the reference experimental structure. 2. **Known mutants and analogs**: reported orthoIL-2 variant termed 3A10 and its cognate orthoIL-2Rβ were included to represent experimentally characterized mutants displaying the desired properties^11,12^. An AF3 model of this complex was also produced alongside its pTM/ipTM values. Additionally, C𝛼 RMSD with respect to the native reference complex was also computed (**Fig. 3c-d**) and showed that 3A10 is in very close structural agreement with both the predicted and actual structure of IL-2–IL-2Rβ*, with RMSDs of 0.562 and 0.219 Å, respectively*. This helped establish a comparative standard to which AF3 predictions of experimentally uncharacterized designs can be compared to AF3 predictions of experimentally characterized variants. 3. **Number of mutations:** At similar ipTM/pTM and RMSD values, a lower number of mutations for a given design is more favorable, the goal is to attain maximum constraint satisfaction with minimum sequence variation. Additionally, a limited sequence variation wrt WT may lower the immunogenic potential of the putative hits at later stages. 4**. Exclude non-canonical binding modes**: RMSD measure for a complex can be biased towards the largest binding partner, we observed some designs with extremely high ipTM/pTM and low RMSDs but still showing non-canonical binding modes (cis and trans binding of *IL-2 and IL-15* at the elbow region of IL-2Rβ were reported in the litterature^41,42^). We filtered these based on a visual inspection of the designed complex structures.

**Figure 3.**
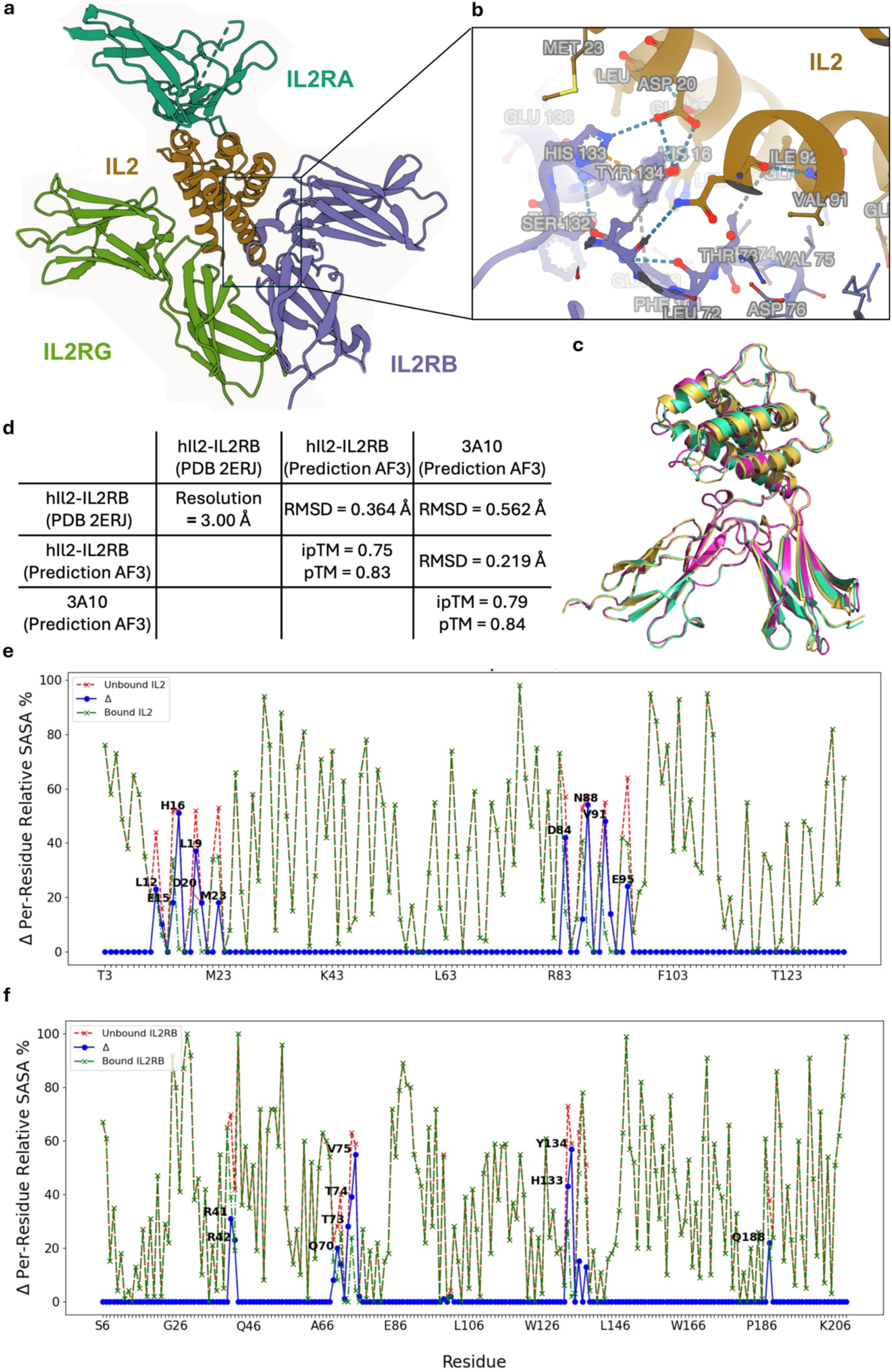
Structural features of the human IL-2R complex interface informs design strategy by unveiling interaction patterns. **a**, Crystal structure of the human quaternary IL-2R signaling complex. IL-2, a 4-bundle α-helical protein of 15 kDa produced by activated T cells, binds to the elbow regions of IL-2Rβ (CD122) and γc (CD132). IL-2Rα (CD25) docks on top of this assembly through IL2 without interacting with the other 2 subunits (PDB: 2ERJ). **b,** Close-up view of the IL-2–IL-2Rβ interface highlighting TYR134 and HIS133 of IL2RB (purple), which form a central interaction hub at the binding surface with IL2 (brown). **c,** Structural alignment of the experimental human (h)IL-2–IL2-Rβ complex (PDB: 2ERJ), an AlphaFold3 (AF3) prediction of (h)IL-2–IL2-Rβ, and the engineered mutant 3A10 predicted by AF3. **d,** Summary table of the structural comparison shown in **c**. Off-diagonal entries report pairwise RMSDs, while diagonal entries report resolution for PDB: 2ERJ and ipTM/pTM scores for AF3 models. **e,** Relative solvent-accessible surface area (rSASA) of IL-2 residues calculated from PDB: 2ERJ. rSASA of IL-2 isolated from IL-2Rβ is shown in red, rSASA of IL2 in the complex is shown in green, and the difference (ΔrSASA) is shown in blue. Residues with a ≥15% decrease in rSASA upon complex formation and an absolute SASA change > 1 Å² are defined as interface residues. **f,** Same rSASA analysis as in **e**, performed for IL-2Rβ.

### Multiple interface definitions expand the accessible design landscape

To explore how alternative definitions of the IL-2–IL-2Rβ binding interface influence the diversity and quality of computational designs, we generated several interface templates for the IL-2–IL-2Rβ system based on distinct criteria. This approach allowed us to probe whether emphasizing different subsets of contact residues could broaden the sequence– structure search space and improve the chances of identifying experimentally viable variants.

Three sets of interfacial residues were defined and recapitulates 7 of the 10 residues previously identified as key contacts^11,12^. **2xSASA interface:** residues showing ≥ 15 % decrease in relative solvent-accessible surface area (rSASA) upon complex formation and an absolute SASA change > 1 Å². This smaller subset (19 residues total) overlaps entirely with the 5 Å set and likewise captures 7 of the 10 previously reported key contact residues (**Fig. 3e-f**). **Reference interface (ref)**: the eight IL-2 residues and two IL-2Rβ positions (His133, Tyr134) explicitly described by Sockolosky et al.^11^ as mediating critical contacts in the *IL-2R* complex (**Fig. 3**). Each interface definition thus represents a distinct, but still overlapping, design template probing how receptor–ligand communication could be rewired without compromising specificity, enabling the exploration of distinct sequence–structure relationships across the interfaces. For each interface subset (5 Å, 2xSASA, ref), we fixed the receptor backbone and designed the interface. The resulting ensembles revealed that interface definition critically shapes interface sequence diversity and mutational patterning across the predicted designs.

### ICPDesign Derived Computational Hits Present Top Quality Metrics

Across the 3 interfaces, the 18 top ranking refined hits showed outstanding ipTM and pTM values, with averages between cognate pairs of 0.724±0.049 and 0.769±0.042, respectively. Displaying metrics close to or within reference WT structure and 3A10 reference orthogonal IL-2–IL-2Rβ mutant (**Fig. 4a-b**). Top designs derived from the 5 Å and 2×SASA sets showed extensive interfacial sequence variation and heterogeneous spatial distribution of predicted stabilizing mutations—reflecting exploration of a broader sequence space and offering a diverse pool of putative orthogonal variants. Interestingly and despite relatively larger sequence variation, 5 Å and 2xSASA in-silico hits showed strong structural similarity to reference 2ERJ (**Fig. 4a-c**). Overall, refined hits comply with secondary constrains on structural deviation from WT IL-2–IL-2Rβ complex measured by an average RMSD of 0.843±0.375 Å (**Fig. 4b**). In contrast, the **ref-based** designs, constrained by fewer positions, yielded two high-quality candidates characterized by minimal sequence variation yet strong compliance with design constraints (**Table 3**). The PCA projection in **Fig. 4c**, shows a group of generated orthoIL-2–IL-2Rβ predicted structures grouped close to the experimental 2ERJ hIL-2–IL-2Rβ, together with the predicted structure of the 3A10 reference mutant. This grouping contained both ref interface hits and four 5 Å interface hits (**Fig. 4a-c**). Initial computational designs of the 5 Å interface yielded a total of 8 candidates with metrics within range (average cognate ipTM = 0.691±0.034, average cognate pTM = 0.739±0.025, non-cognate ipTM average non-cognate ipTM = 0.243±0.096) (**Table 3**). Estimating the RMSDs with respect to the reference WT, these models showed substantial deviation from the WT structure (RMSD >> 1 Å, average RMSD = 4.001±2.085 Å). 1 out of the 8 showed an RMSD of 1.378 Å (**Suppl Table 2**). With these first set of encouraging results, we ran a smaller second sampling round that yielded 5 top quality hits, one of which was discarded for showing RMSD to WT = 5.1 Å, mainly driven by deviations in the mutant IL-2Rβ subunit **(Suppl Table 3)**. The 4 remaining designs showed outstanding quality metrics (**Table 3**), with average cognate ipTM = 0.745±0.076, average cognate pTM 0.765±0.062, average RMSD to WT = 1.057±0.356 Å and average non-cognate ipTM = 0.178±0.0789, suggesting strong binding ability for cognates, high structural quality of predictions, a close structural agreement between designs and WT IL-2–IL-2Rβ complex, and orthogonal properties wrt to WT IL-2–IL-2Rβ. Furthermore, the interface composition of these designs was free of deletions. However, designs based on the 5 Å interface all showed extensive mutations on the templates, with nearly all 27 interfacial positions being mutated for all hits. The 2xSASA interface on the other hand, yielded 12 candidates before applying secondary constraints with average cognate ipTM = 0.718±0.036, average cognate pTM = 0.759±0.05 and average RMSD to WT = 1.093±1.425 Å (**Suppl Table 4**). Applying secondary constraints, 1 hit was discarded because of its large RMSD to WT (6.1 Å), leaving 11 designs with ideal metrics (**Table 3**). 2xSASA results, showed lower numbers of mutations on the templates ∊ [14-19] compared to the 5 Å interface, and average non-cognate ipTM = 0.14±0.052, suggesting a strong orthogonality potential for these top-ranking hits with a lower mutational burden. Finally, sampling the Ref interface, we found 2 high quality hits which satisfied all secondary constraints, a substantially smaller set when compared to the 2 other interfaces (**Suppl Table 5**). One of the 2 hits presented deletions at the interface, the other one showed ideal parameters. This hit named 69R3, showed cognate ipTM = 0.77 and pTM = 0.84 within references range (WT IL-2–IL-2Rβ and 3A10 *ortho*IL-2–IL-2Rβ, ***Fig. 3d***), [0.75, 0.79] and [0.83, 0.84], respectively, suggesting a strong potential for high binding affinity for cognates and overall, very high structural confidence of the predicted model (**Suppl Table 1**). Additionally, this hit presented a conformation in very close agreement with the reference WT IL-2–IL-2Rβ complex, showing a staggering **0.349 Å** C𝛼 RMSD wrt WT IL-2–IL-2Rβ, the lowest observed across all interfaces (**Suppl Table 1**), with a binding mode quasi-identical to WT (**Fig. 4a-b**). Moreover, a non-cognate ipTM value of 0.49 suggests that the designed binder is orthogonal to the WT receptor. 69R3 only presents 7 mutations, 6 on IL-2 and only 1 mutation on IL-2Rβ, where the reference 3A10 shows 10 mutations on the WT template with 8 on IL-2 and 2 on IL-2Rβ, satisfying all design constraints with minimal sequence variation. Finally, we sought to computationally investigate how the predicted structural variations induced by interface mutation influenced the dynamic of the generated putative orthoIL-2–IL-2Rβ. The Gaussian Network Model (GNM), a simple yet powerful model describing the residue-agnostic dynamic behavior of proteins in crystals, is used to assess and compare the intrinsic dynamic encoded in protein structure coordinates^43–45^.

**Figure 4.**
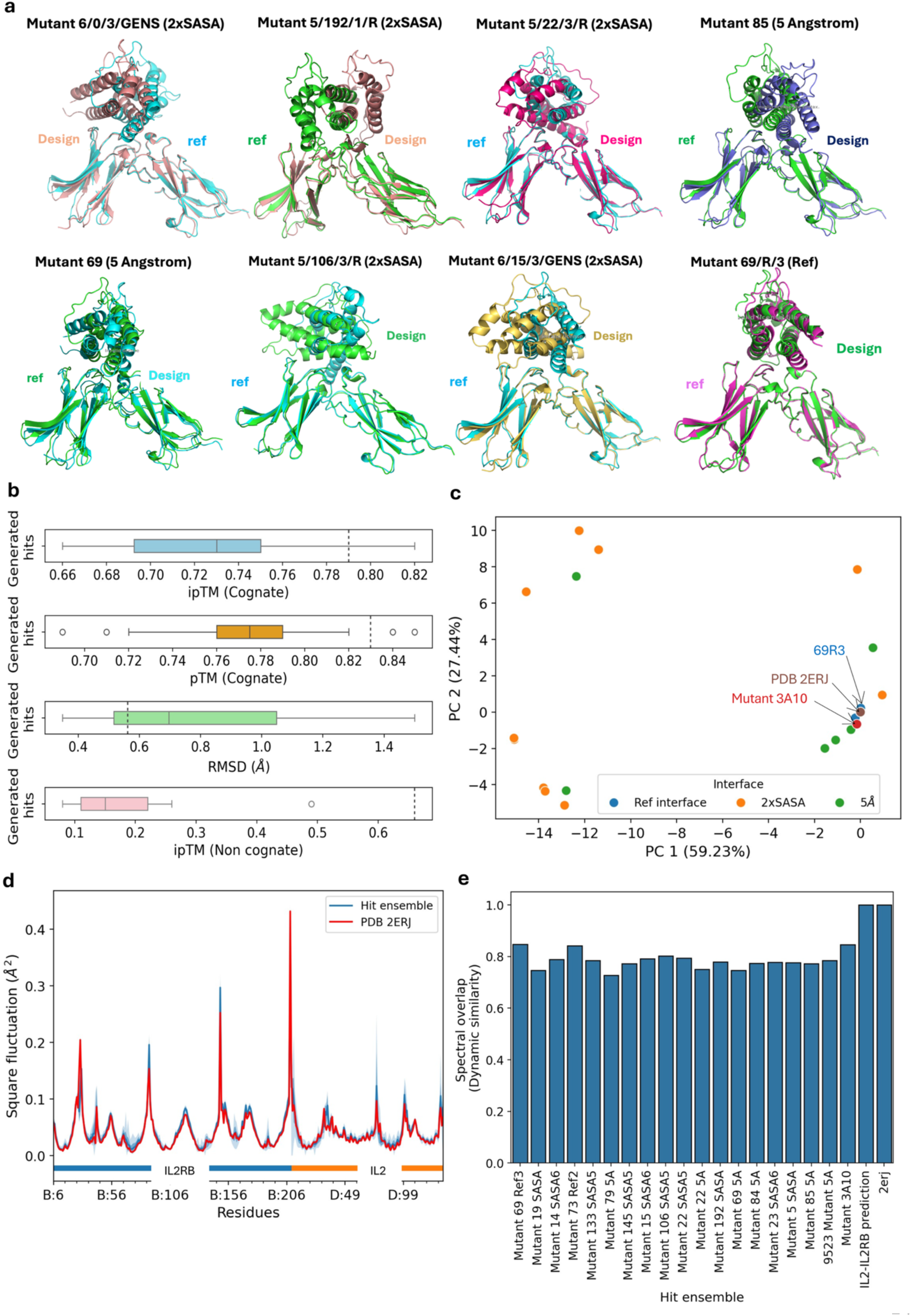
Top-ranked IL-2–IL-2Rβ *in-silico* hits closely match native structure and motion. **a**, Representative structural alignments of the top eight ranked mutant hits from each engineered interface (labelled design), superposed onto the reference human (h)IL-2–IL-2Rβ complex (PDB: 2ERJ; labelled ref). **b,** Box plots summarizing structural and confidence metrics of the hit ensemble. From top to bottom, distributions of ipTM and pTM of AF3 models of hits, dotted lines indicate reference values: ipTM and pTM obtained from an AF3 model of the WT (h)IL-2–IL-2Rβ complex, distribution of RMSD (Å) of hits to reference PDB: 2ERJ, dotted line represent RMSD of the 3A10 mutant relative to PDB: 2ERJ. The final box plot, describes the distribution of ipTM of mutant IL-2 hits in complex with (h)IL-2Rβ, the dotted line represent the ipTM value recovered from a model of AF3 model of orthogonal IL-2 (3A10) bound to (h)IL-2Rβ. **c,** Principal component analysis (PCA) of Cα atoms from a structural ensemble comprising all mutant hits, PDB: 2ERJ, and the 3A10 mutant. The percentage of variance explained by each principal component is reported on the axes. **d,** Square fluctuations of Cα atoms (Å^2^) calculated by a Gaussian network model (GNM). Fluctuations derived from the ensemble of hits are shown in blue and compared to the reference structure (PDB: 2ERJ) shown in red, reflecting expected positional variability due to intrinsic, coordinate-encoded motions. **e,** Root mean square inner product (RMSIP) analysis comparing the 20 dominant collective modes of motion. One-by-one comparisons are shown for each hit, the predicted reference mutant 3A10, and the hit ensemble, all evaluated with respect to PDB: 2ERJ, for which RMSIP = 1. 3A10 and the mutant 69R3 exhibit the highest RMSIP values, indicating the greatest similarity of collective motions to the reference.

**Table 3.** Summary of the computational hits obtained after applying secondary constraints. This second round of filtration halved the number of hits. cog; cognate, non-cog; non-cognate. SD: Standard deviation. Interfaces are defined in **Table 2**.

| Model | Interface | # of refined hits | Average ipTM cog ± SD | Average pTM cog ± SD | Average RMSD to WT (Å) ± SD | Average ipTM non-cog ± SD |
| --- | --- | --- | --- | --- | --- | --- |
| 1 | 5 Å | 1 | 0.660 | 0.720 | 1.378 | 0.190 |
| 2 | 5 Å | 4 | 0.745±0.076 | 0.765±0.062 | 1.057±0.356 | 0.178±0.079 |
| 3 | 2xSASA | 11 | 0.716±0.036 | 0.750±0.050 | 0.829±0.410 | 0.140±0.052 |
| 5 | Ref | 2 | 0.720±0.050 | 0.815±0.025 | 0.355±0.006 | 0.490±0.000 |

**Fig. 4c** displays the mean square fluctuation (MSF, Å^2^) of C-alpha atoms around their predicted or experimental spatial positions, predicted by a GNM model for the ensemble of hit structures (blue curve) described in **Suppl Table 1**. The red line depicts the MSF predicted by a GNM model for the reference 2ERJ. Overall, the ensemble’s average MSF is highly similar to the reference, with curves mostly overlapping, suggesting a conserved collective dynamic despite limited sequence and structure variation at the interface. A one-by-one investigation of the hits predicted GNM modes of motion shown in **Fig. 4d**, reveals a high degree of dynamic similarity between the reference 2ERJ, the predicted reference mutant 3A10 and the hit ensemble. 3A10 and 69R3 show the highest Root Mean Square Inner Product (RWSIP), essentially a similarity measure of collective modes of motion ranging between 0 (no similarity) and 1 (identical)^46^, to the reference 2ERJ indicating a high dynamic fidelity to the WT.

## Discussion

This study is a demonstration of in-silico generation of prioritized sets of orthoIL-2–IL-2Rβ pairs. Though the focus of the results was on a restricted set of high scoring mutants, with designed variant 69R3 presenting remarkable quality metrics with minimal sequence variation, the experimentally preferred strategy would aim at testing few thousands of ICPDesign generated orthoIL-2–IL-2Rβ pairs on one or more interface. Investigated high quality in-silico hits on the other hand, offer a structural glimpse of the best predicted mutants. These in-silico hits were generated by computationally exploring a fraction of the experimental sequence space generated by Sockolosky et al. Ideally, one would translate the computational exploration into more focused test sets for the experimental assessment of the biological properties of model systems expressing these proteins. An ideal subsequent study would reproduce the experimental strategy established by Sockolosky et al. at a smaller scale by evaluating few thousands (∼10^3^) of ICPDesign generated orthoIL-2-IL-2Rβ mutants.

A major strength of the approach is its grounding in structure and experimental gold standards, with interface predictions informed by state-of-the-art AlphaFold3. The rational choice of metrics and interface definitions, together with the strong theoretical basis of the Potts model, provides an interpretable, context-specific, dataset (cMSA) adjustable statistical model. With its original decomposition-sampling approach, the current interpretation of the CSG model is that it samples expected interaction interfaces, thus the need for exploring the exceptionally high probability ends of the likelihood distribution for putative hit-enriched subsets. Due to its unsupervised nature, the model only requires knowledge of truly interacting pairs to build the concatenated alignment, without explicit knowledge of negative interaction. In practice, relatively few (20-50) independent CSG replicas, compared to previous attempt by Malinverni and Babu^20^ (>180 MCMC replicas), were sufficient to identify top in silico candidates. Moreover, only a few thousands of top-ranking mutants, across interface sets, were structurally modelled with AF3, supporting computational efficiency. Expanding the number of CSG replicas exploring the sequence landscape could further refine the model’s parameter set and performances. Structure prediction currently represents a bottleneck due to reliance on AlphaFold3; by only exploring a few thousands of prioritized sequences, the full power of in-silico screening is not attained. Input MSA retrieval using the AF3 pipeline do consume a significant amount of time and hinders perspective of large scale in-silico prediction. The “single-sequence mode” of AF3 may present an alternative by performing prediction with no input MSA, cutting processing time. However, Peng et al. did observe a decrease of 20% of AF3’s structural accuracy when used in this mode *versus* with input MSA^39^. While the implementation shows early but genuine generative behaviour beyond uniform sequence sampling, the generative mode of the CSG framework tends to converge toward restricted subsets of a few thousands of sequences, limiting the scope and diversity of generated candidate variants. In addition, the training cMSA is relatively small when compared to Malinverni and Babu’s, with the important difference that the authors studied a bacterial system with many known homologs^20^. Additionally, the feature set based on statistical coupling may limit the diversity and nature of learned interactions. Expanding the training data (i.e increasing the number of interaction examples) would increase the power of the model, with the latter’s parameter set size varying quadratically with the size of the mutation interface (**Methods**).

Within the current literature, orthogonal IL-2–activated CAR T cell systems have shown encouraging pre-clinical results, with studies reporting selective and controlled expansion, superior engraftment, potent tumor suppression, and reduced off-target immune activation despite toxicities at higher doses^11,12^. Beyond IL-2, closely related cytokine systems such as IL-15 that also signals through the IL-2Rβ subunit, as well as distinct pathways involving TNF or IFNγ, could be explored to systematically determine which signalling axis best optimizes pre-clinical parameters.

## Methods

### Constrained Sequence Generator (CSG)

The Constrained Sequence Generator (CSG) is a generative model designed to produce novel protein sequences, with the aim to satisfy user-defined structural or functional constraints (**Suppl Fig. 1b**). Unlike earlier Potts-based frameworks focused only on scoring sequences, CSG can actively generate variants that align with evolutionary and biophysical data. At its core, CSG builds on the Potts model, which captures co-evolutionary patterns between amino acid positions in related proteins. Each sequence is assigned an energy value, or “Hamiltonian” (**Suppl Equation 1-2**). The “Hamiltonian”, a legacy nomenclature from the Potts model, is a scoring function used to evaluate sequences (**Suppl Fig. 1b**). The score is a linear combination of stochastic parameters ({𝞱}, {ɸ}) to be optimized, and statistics extracted from concatenated Multiple Sequence Alignments (cMSA). This quantity is then used to measure a probability density for a given sequence using the Boltzmann distribution (**Suppl Equation 3-5**). CSG leverages the interface probability decomposition property described in (**Suppl Equation 1-5**), to treat each mutation position, within a protein sequence, independently. As such, each mutation position composition is modelled by a random probability vector over the 20 natural amino acids and a gap state (i.e. deletion) (**Suppl Equation 6**).

Due to the positional independence, this vector is drawn from a Dirichlet distribution, parametrized with the sequence probability density in (**Suppl Equation 8**). This configuration allows the Dirichlet prior probability to reflect the value of the Boltzmann probability of the sequence. The Potts model framework, however, doesn’t allow for direct sampling using the Boltzmann distribution, due to the impossibility to calculate a normalization constant (**Suppl Equation 7**). Most authors used random sampling to optimize likelihood estimators and later randomly explore and score the sequence space. To introduce non-random generative capabilities, we used the Dirichlet probability vectors per position as input to a multinomial distribution (**Suppl Equation 9**). This process allows to build a prior on the probability of a sequence given the data and the imposed constraints, the well characterized Dirichlet-Multinomial distribution. From this prior one can now draw sequences which likelihood is assessed with the Boltzmann distribution, yielding a posterior probability termed Boltzmann-Dirichlet-Multinomial (BDM) (**Suppl Equation 10-11**). Additional steps described in (**Suppl Equation 12-13**), yield the evidence lower bound (ELBO), an estimator incorporating information from the prior and the likelihood. Maximizing the ELBO is equivalent to minimizing the KL divergence between the generative prior and the likelihood estimator, resulting in an increasingly relevant sampling by making the generative model iteratively closer to the Boltzmann distribution^24,47,48^. This model is, to the best of our knowledge, a novel simulator of the Boltzmann distribution incorporating the affinity/orthogonality constraints by construction, and capable of sampling relevant parts of the sequence space. To enforce constraints in a systematic way, the “Hamiltonian” of the generated and the training sequences are compared to that of a set of random sequences, (**Suppl Equation 14**). This comparison allows to penalize models which generate sequences less likely than a random generator and/or deems actual training sequences likewise. The orthogonality constraint is introduced by calculating the “Hamiltonian” of a pair constituted by generated sequences and the constraint sequence (**Suppl Fig. 1c**). Using the constrained “Hamiltonian” and adding to it the L2 norm of the model, we obtain the constrained and regularized expected log likelihood of the model (**Suppl Equation 15-17**). This quantity is then used to build the objective function of the model, which is used to assess proposal parameters using the Metropolis-Hastings criteria (**Suppl Equation 18-20**). The last piece of the CSG algorithm is the optimization of the model’s parameters {𝞱} and {ɸ} used to compute the score “Hamiltonian”. Given the large set of parameters of the model, we opted for the Hamiltonian Monte Carlo (HMC), an efficient optimization algorithm capable of breaking away from the local behavior of the RW, its large rejection rate and burdensome computations. HMC exploit properties of Hamiltonian Flow by proposing bolder moves in state space while displaying higher acceptance rate, and scales better to high-dimensional problems^47,49^.Though originating in physics, HMC can be applied to most problems with continuous state spaces. The first step is to define a “Hamiltonian” function in terms of the log probability distribution we wish to sample from. A stochastic perturbation is introduced by an auxiliary “momentum” variable, which typically has an independent Gaussian distribution (**Suppl Equation 21-22**). HMC alternates updates for these variables in which a new state is proposed by integrating Hamilton’s equations of motion using the leapfrog method (**Suppl Equation 23-27**). Proposed states can be distant from the current state but nevertheless have a high probability of acceptance^47,49^. In summary (**Suppl Fig. 1**), CSG is a stochastic process composed of 2 main compartments. First HMC produces proposal parameters from the data, that are then used by the BDM process to generate sequences. By iteratively optimizing the set of proposed parameters the model becomes able to sample regions of the sequence space increasingly likely to satisfy the imposed constraints. The model is distributed in the form of a Python package (See **Data availability**).

### Model training and datasets

#### Dataset

We trained the CSG algorithm on a dataset of homologs of the *IL-2–IL-2Rβ* system. The Dataset consisted of a Multiple Sequence Alignment (MSA) containing 527 pairs of concatenated IL-2–IL-2R*β* pairs across different species (**Suppl Fig. 1b**) downloaded from Uniprot and based on two independent pBlast of canonical human *IL-2* and *IL-2Rβ*, using default parameters^50^. The *IL-2* and IL-2R*β* retrieved from the pBlast yielded 262 and 922 sequences, respectively. The two sets of sequences were then independently aligned using MAFFT version 7 with default parameters^51^. Prior to the specie-wise concatenation, the 2 MSAs were filtered using the refineMSA function of the ProDy suite^52^. All the sequences in the alignments were trimmed to the size of the canonical Human IL-2 and IL-2R*β* sequences, only sequences displaying a row occupancy ≥0.8 were kept. To reduce redundancy and sampling bias due to overrepresented organisms, we retained sequences pairs having a maximum sequence similarity of 98%. Cognate *IL-2–IL-2Rβ* were then matched for each organism and concatenated. We assumed that all isoforms bind to their cognates for a given organism^20^. Residues not included in the experimentally resolved 2ERJ structure were filtered out yielding an MSA of 571 paired *IL-2–IL-2Rβ* sequences consisting of 343 columns (residues). **Fig. 5 a-c** recapitulates the characteristics of the training set.

**Fig. 5.**
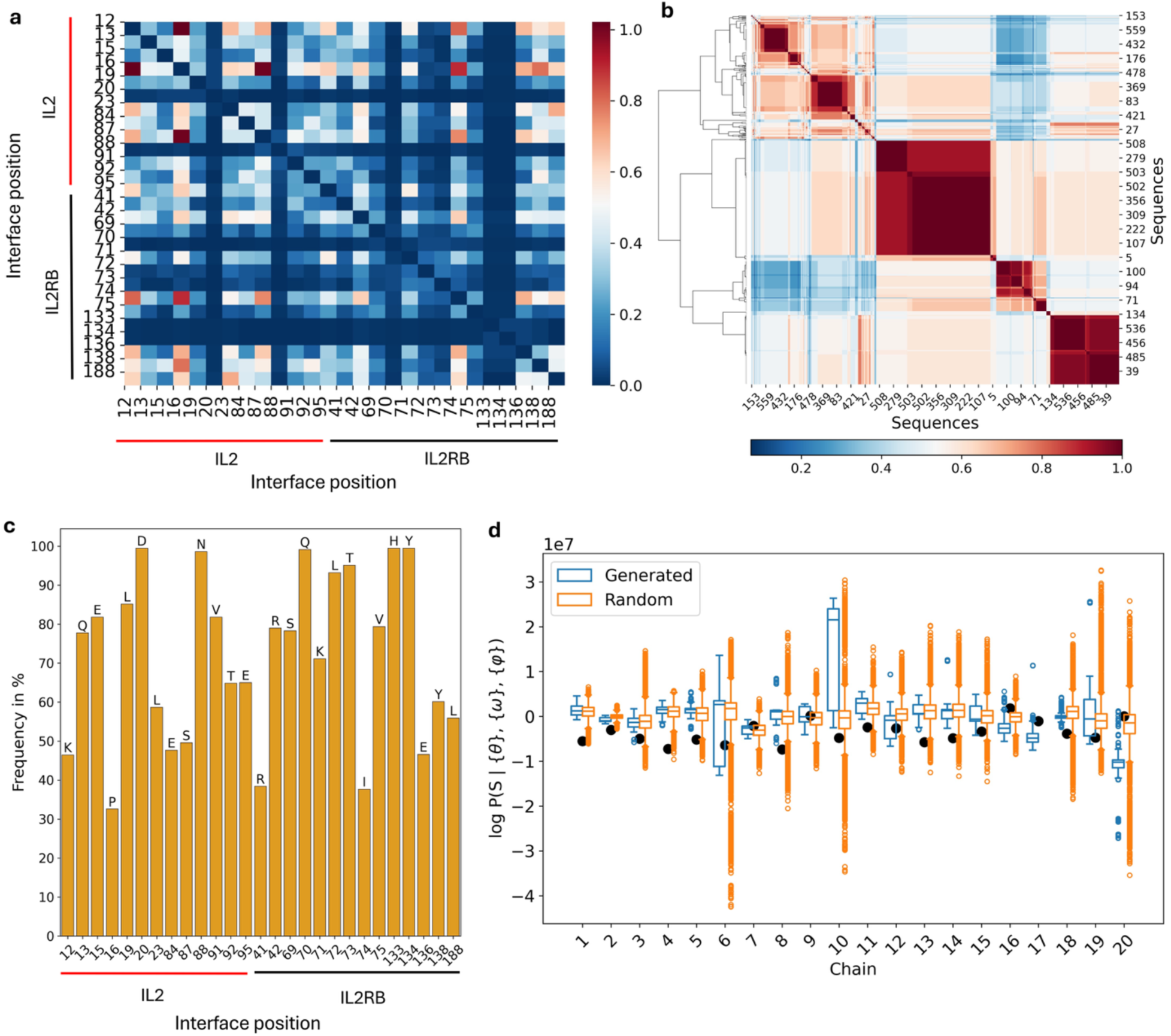
Evolutionary signals guide orthogonal IL-2–IL-2Rβ interface design. **a**, Mutual information matrix computed over residues within 5 Å at the IL-2–IL-2Rβ interface from the concatenated multiple sequence alignment (cMSA). Red–orange signals denote strong intra- and intermolecular residue associations, whereas deep blue horizontal bands indicate highly conserved positions. **b,** Pairwise sequence identity matrix of the IL-2–IL-2Rβ interface cMSA with hierarchical clustering (Ward variance minimization, Euclidean distance) shown as a row dendrogram. Clustering reveals three major sequence groups. **c,** Residue frequency distribution across IL-2–IL-2Rβ interface positions. The x axis represents positions within the 5 Å interface and the y axis shows residue frequency. Bars are labeled with the most frequent residue at each site. Four positions show near-complete conservation: H133, Y134 and Q70 in IL-2Rβ, and N88 and D20 in IL-2. **d,** Log-likelihood distributions of sequences generated by 20 independent Markov chains used to train model 1 (**Table 1**). Each chain generated 10,000 sequences that were scored under the learned model (orange box plots) and compared to randomly generated sequences (uniform distribution, orange box plots) evaluated under the same likelihood function. Chain 10 displays a marked shift toward higher likelihood values. The top 1.35% highest scoring yielded the hits reported for model 1, all were from chain 10. Y axis scale 10^7^.

#### CSG training

A typical CSG model is trained using a multi-replica strategy, where W independent models perform optimization and take a concatenated MSA as input. Each replica requires the user to define the positions {q} on which the model will operate to generate variants (i.e. interface), and the 7 hyperparameters of the model (**Supplementary**). Additionally, users can define a constrained binder by indicating its position in the cMSA. To optimize the Potts model parameters, each replica runs M HMC trajectory consisting in L leapfrog integration step at resolution Δt. Each trajectory yields a set of proposal parameters for BDM sampling as described previously and summarized in (**Suppl Fig. 1a**). The number of parameters (S) of the model is given by the sum of the dimensions of the frequency matrix F and the coupling tensor C^2^, S depends directly on the number of variable positions q:

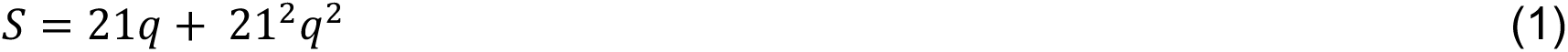

Model training yields W sets of optimized parameters that can be used for data generation with the BDM. A single BDM process can draw Q training samples composed of N sequences. 3 sampling strategies can be used to generate data with the optimized parameters (**Table 1**): 1. Multi-replica sampling: each replica generates N sequences using its own optimized parameters, sampling results are pooled across predictors, their replica likelihood is calculated, and sequences are ranked (**Fig. 5d**). 2. Ensemble sampling: Parameters are averaged across replicas and used to draw samples from a BDM with ensemble parameters, sequences ensemble likelihood is calculated followed by ranking. 3. Random sampling: Random sequences are drawn from a uniform distribution (i.e. independent equiprobability of all residues at all {q} positions), likelihoods are calculated and sequences are ranked. User is encouraged to diversify sampling strategies to increase hit rate. CSG uses a Pytorch v2.9.1 implementation of the HMC-BDM duo, allowing for cuda accelerated model training, sequence sampling and likelihood estimation.

### Structural Modelling

#### AlphaFold3 pipeline

AlphaFold3 was installed following the official source code repository guidelines and model parameters were obtained under the required terms of use. Input definitions were prepared in JSON format with chain identifiers and IL-2 and IL-2Rβ sequences. Each job included multiple random seeds to generate an ensemble of structural predictions. Predictions were executed on GPU-equipped compute nodes (Nvidia A100, Nvidia l40s). Output files included coordinate files (.cif) and confidence metrics such as the pTM/ipTM and other model scores for assessing prediction reliability. The software was used in MSA mode, where each prediction automatically retrieves an MSA^39^. AlphaFold3’s data pipeline runs Jackhmmer searches against multiple sequence databases (UniRef90, MgNify, UniProt..) to find homologous sequences and construct the alignment. Results from different databases are merged, deduplicated, and formatted into unpaired and paired MSAs for use by the model^53^.

#### Structural analysis

Structural visualization was prepared using PyMOL v3.0.3 and Mol* (Mol* Viewer)^54^. Structural alignments and root-mean-square deviation (RMSD) calculations were carried out in PyMOL using the align command, all reported RMSD values were computed after optimal C𝛼 structural alignment. Solvent-accessible surface area (SASA) calculations were performed in PyMOL using the get_area and get_sasa_relative commands for the absolute SASA and relative SASA, respectively. Distance-based 5 Å interface was identified in PyMOL by selecting residues within a 5 Å distance cutoff between chains. Ensemble-level structural and dynamical analyses were carried out using ProDy^52^. A C𝛼 protein ensemble was constructed using buildPDBEnsemble, including all in-silico predicted structures together with the experimental reference structure PDB: 2ERJ, which served as the alignment reference for all ensemble members. Covariance matrix was computed from the aligned ensemble, followed by principal component analysis (PCA). Projections of individual structures onto principal components were obtained using ProDy’s PCA projection utilities. Collective dynamics of the ensemble were further analyzed using the Gaussian Network Model (GNM) as implemented in ProDy, allowing the assessment of dominant intrinsic modes of motions across the structural ensemble.

### Data Availability

The concatenated multiple sequence alignment (cMSA) used to train CSG is publicly available at GitHub (https://github.com/ghostface16/ICPDesign/tree/main), and is the only input file to the model. Structural files and interface composition data supporting this study are available from the corresponding author upon reasonable request.

### Code Availability

CSG is distributed as a Python package and is available at GitHub (https://github.com/ghostface16/ICPDesign/tree/main). The repository also includes a detailed Jupyter notebook that guides users through the constrained protein design workflow. Use of the provided workflow requires access to a local installation of AlphaFold3 (https://github.com/google-deepmind/alphafold3).

