## Supplementary for "Rational Design of Selective IL-2-based Activators for CAR T Cells Using AlphaFold3 and Physics-Informed Machine Learning"

**Supplementary Table 1. Summary of high-quality *in-silico* hits with estimates of primary and secondary constraints metrics.** IL-2Rβ–IL-2*: denotes the predicted complex formed by WT IL-2Rβ and mutant IL-2*.

| Mutant | ipTM | pTM | RMSD wrt PDB: 2ERJ (Å) | Interface | # mutations | IL-2Rβ–IL-2* ipTM |
|---|---|---|---|---|---|---|
| 6/0/3/GENS | **0.73** | **0.78** | **0.667** | 2xSASA | 16 | **0.16** |
| 5/22/3/R | **0.79** | **0.78** | **0.726** | 2xSASA | 14 | **0.15** |
| 5/192/1/R | **0.76** | **0.8** | **0.746** | 2xSASA | 17 | **0.15** |
| 5/5/2/G | 0.68 | 0.69 | 1.089 | 2xSASA | 19 | **0.25** |
| 6/14/3/GENS | **0.70** | **0.77** | **0.527** | 2xSASA | 17 | **0.08** |
| 6/15/3/GENS | **0.73** | **0.76** | **0.452** | 2xSASA | 17 | **0.1** |
| 5/106/3/R | **0.7** | **0.77** | **0.414** | 2xSASA | 14 | **0.11** |
| 5/133/3/R | **0.72** | **0.77** | 1.247 | 2xSASA | 15 | **0.09** |
| 5/145/3/R | 0.68 | **0.76** | **0.85** | 2xSASA | 15 | **0.19** |
| 6/23/1/R | **0.73** | **0.79** | 1.304 | 2xSASA | 17 | **0.16** |
| 69/R/3 | **0.77** | **0.84** | **0.349** | Ref | **7** | **0.49** |
| 22 | 0.67 | **0.71** | 1.5 | 5 Å | 22 | **0.11** |
| 69 | **0.8** | **0.73** | 1.145 | 5 Å | 23 | **0.11** |
| 84 | 0.69 | **0.77** | **0.657** | 5 Å | 26 | **0.23** |
| 85 | **0.82** | **0.85** | **0.927** | 5 Å | 25 | **0.26** |
| 9523 | 0.66 | **0.72** | 1.378 | 5 Å | 25 | **0.19** |
| 73/R/2 | 0.67 | **0.79** | **0.36** | Ref | **7** | **0.49** |
| Ref mutant 3A10 (prediction) | 0.79 | 0.84 | 0.562 | Ref | 10 | 0.66 |
| Canonical hIL-2–IL-2Rβ (prediction) | 0.75 | 0.83 | 0.364 | Canonical | / | / |

**Supplementary Table 2. Summary of initial *in-silico* hits obtained for the 5Å interface, with estimates of primary and secondary constraints metrics.** Despite high structural quality and interface metrics, all hits show substantial structural deviation when compared to the experimental reference, reflected by high RMSD values. IL-2Rβ–IL-2*: denotes the predicted complex formed by WT IL-2Rβ and mutant IL-2*.

| Mutant | ipTM | pTM | RMSD wrt PDB: 2ERJ (Å) | IL-2Rβ–IL-2* ipTM | # mutations |
|---|---|---|---|---|---|
| 5890 | 0.65 | **0.71** | 3.616 | **0.44** | 23 |
| 5963 | **0.74** | **0.78** | 6.256 | **0.32** | 23 |
| 5989 | **0.7** | **0.74** | 6.358 | **0.17** | 22 |
| 6360 | 0.65 | **0.71** | 6.472 | **0.16** | 23 |
| 3219 | **0.71** | **0.75** | 2.173 | **0.26** | 23 |
| 2845 | 0.66 | 0.62 | 6.285 | **0.16** | 23 |
| 1114 | 0.66 | 0.62 | 5.281 | **0.3** | 23 |
| 1269 | **0.7** | **0.74** | 3.501 | **0.22** | 23 |
| 9587 | **0.72** | **0.76** | 2.255 | **0.18** | 23 |
| 9523 | 0.66 | **0.72** | **1.378** | **0.19** | 25 |
| Ref mutant 3A10 (prediction) | **0.79** | **0.84** | **0.562** | 0.66 | 10 |
| Canonical hIL-2–IL-2Rβ (prediction) | **0.75** | **0.83** | **0.364** | / | / |

**Supplementary Table 3. Summary of subsequent *in-silico* hits obtained for the 5Å interface, with estimates of primary and secondary constraints metrics.** This model yielded candidates with high structural quality and interface metrics, four out of five hits show close structural fidelity to the experimental reference, reflected by RMSDs ~1Å. IL-2Rβ–IL-2*: denotes the predicted complex formed by WT IL-2Rβ and mutant IL-2*.

| Mutant | ipTM | pTM | RMSD wrt PDB: 2ERJ (Å) | IL-2Rβ–IL-2* ipTM | # mutations |
|---|---|---|---|---|---|
| 22 | **0.67** | **0.71** | 1.5 | **0.11** | 22 |
| 69 | **0.8** | **0.73** | 1.145 | **0.11** | 23 |
| 79 | **0.78** | **0.74** | 5.1 | **0.12** | 24 |
| 84 | **0.69** | **0.77** | **0.657** | **0.23** | 26 |
| 85 | **0.82** | **0.85** | **0.927** | **0.26** | 25 |
| Ref mutant 3A10 (prediction) | 0.79 | 0.84 | 0.562 | 0.66 | 10 |
| Canonical hIL-2–IL-2Rβ (prediction) | 0.75 | 0.83 | 0.364 | / | / |

**Supplementary Table 4. Summary of *in-silico* hits obtained for the 2xSASA interface, with estimates of primary and secondary constraints metrics.** This model yielded candidates with high structural quality and interface metrics, four out of five hits show close structural fidelity to the experimental reference, reflected by RMSDs ~1Å. IL-2Rβ–IL-2*: denotes the predicted complex formed by WT IL-2Rβ and mutant IL-2*.

| Mutant | ipTM | pTM | RMSD wrt PDB: 2ERJ (Å) | # mutations | IL-2Rβ–IL-2* ipTM |
|---|---|---|---|---|---|
| 6/0/3/GENS | **0.73** | **0.78** | **0.667** | 16 | **0.16** |
| 5/22/3/R | **0.79** | **0.78** | **0.726** | 14 | **0.15** |
| 5/192/1/R | **0.76** | **0.8** | **0.746** | 17 | **0.15** |
| 5/5/2/G | **0.68** | **0.69** | 1.089 | 19 | **0.25** |
| 5/19/1/R | **0.67** | 0.62 | 1.107 | 19 | **0.12** |
| 6/14/3/GENS | **0.70** | **0.77** | **0.527** | 17 | **0.08** |
| 6/15/3/GENS | **0.73** | **0.76** | **0.452** | 17 | **0.1** |
| 5/106/3/R | **0.7** | **0.77** | **0.414** | 14 | **0.11** |
| 5/133/3/R | **0.72** | **0.77** | 1.247 | 15 | **0.09** |
| 5/145/3/R | **0.68** | **0.76** | **0.85** | 15 | **0.19** |
| 6/23/1/R | **0.73** | **0.79** | 1.304 | 17 | **0.16** |
| 6/44/1/R | **0.65** | **0.68** | 6.514 | 15 | **0.13** |
| Ref mutant 3A10 (prediction) | 0.79 | 0.84 | 0.562 | 10 | 0.66 |
| Canonical hIL-2–IL-2Rβ (prediction) | 0.75 | 0.83 | 0.364 | / | / |

**Supplementary Table 5. Summary of *in-silico* hits obtained for the Ref interface, with estimates of primary and secondary constraints metrics.** This model yielded the least amount of hits, however, mutant 69R3 derived from this model, present ideal characteristic metrics. IL-2Rβ–IL-2*: denotes the predicted complex formed by WT IL-2Rβ and mutant IL-2*.

| Mutant | ipTM | pTM | RMSD wrt PDB: 2ERJ (Å) | # mutations | IL-2Rβ–IL-2* ipTM |
|---|---|---|---|---|---|
| 69/R/3 | **0.77** | **0.84** | **0.349** | **7** | **0.49** |
| 46/R/3 | 0.6 | **0.71** | **0.682** | **8** | **0.14** |
| 73/R/2 | **0.67** | **0.79** | **0.36** | **7** | **0.49** |
| Ref mutant 3A10 (prediction) | 0.79 | 0.84 | 0.562 | 10 | 0.66 |
| Canonical hIL-2–IL-2Rβ (prediction) | 0.75 | 0.83 | 0.364 | / | / |

**Constrained Sequence generator**

Decomposed score H: per residue $k_1$ observed at position j:

$$H_{jk}(S_{jk} \mid F,\ C^2, \{\theta\}, \{\phi\}) = \theta_{jk_1} F_{jk_1} + \sum_{i=1}^{q} \sum_{k_2=1}^{21} \varphi_{ijk_1k_2} C^2_{ijk_1k_2} + Constraints \tag{1}$$

Total score H of a sequence:

$$H(S \mid F,\ C^2, \{\theta\},\ \{\phi\}) = \sum_{j=1}^{q} \theta_{jk_1} F_{jk_1} + \sum_{i=1}^{q} \sum_{j=1}^{q} \sum_{k_2=1}^{21} C^2_{ijk_1k_2} + Constraints \tag{2}$$

Using the exponential property, we show that the probability to pick a residue $k_1$ at position j can be decomposed as follows:

$$\boldsymbol{P(S_{jk} \mid \{\theta\},\ \{\phi\},\ data) \propto exp \left\{ {-H_{jk}(S_{jk} \mid \{\theta\}, \{\phi\},\ data)} \big/ {T} \right\}} \tag{3}$$

Now, the probability of an amino acid sequence can be expressed as:

$$P(S \mid \{\theta\},\ \{\phi\},\ data) \propto \prod_{j=1}^{q} \boldsymbol{exp} \left\{ {-\boldsymbol{H_{jk}(S_{jk} \mid \{\theta\},\ \{\phi\},\ data)}} \big/ {\boldsymbol{T}} \right\} \tag{4}$$

Using simple exponential property, (4) gives a formulation equivalent to a Potts model:

$$P(S \mid \{\theta\},\ \{\phi\},\ data) \propto exp \left\{ {-\sum_{j=1}^{q} H_{jk}(S_{jk} \mid \{\theta\}, \{\phi\},\ data)} \big/ {T} \right\} \tag{5}$$

Assigning each possible residue k at each mutable position j the Boltzmann density $\propto P(S_{jk} \mid \{\theta\},\ \{\phi\},\ data)$ such that:

G = (6)

| | 1 | .. j .. | q |
|---|---|---|---|
| 1 | $\propto P(S_{1,1} \mid \{\theta\},\ \{\phi\},\ data)$ | .. | $\propto P(S_{1,q} \mid \{\theta\},\ \{\phi\},\ data)$ |
| ..k.. | ⋮ | . | ⋮ |
| 21 | $\propto P(S_{21,1} \mid \{\theta\},\ \{\phi\},\ data)$ | .. | $\propto P(S_{21,q} \mid \{\theta\},\ \{\phi\},\ data)$ |

The columns of G do not satisfy Kolmogorov's axioms:

$$1 \geq G_{ij} \geq 0 \quad ; \quad \sum_{i=1}^{21} G^T_{ij} = 1 \tag{7}$$

To Satisfy Equation (7), V is derived from G, such that:

$$V_j^T = p(S_j) \sim Dir(G_{1,j} \ldots G_{21,j}) \quad (8)$$

Draw sequences position-wise using a multinomial distribution:

$$(S_{j,1} \ldots S_{j,N}) | V_j^T \sim Mult(N, p(S_j)) \quad (9)$$

Posterior Boltzmann-Dirichlet-Multinomial (BDM):

$$p(data, \{\theta\}, \{\varphi\}|S) \propto p(S \mid data, \{\theta\}, \{\varphi\}) \quad p(S) \quad (10)$$

If N independent sequences are drawn, Equation 10 becomes:

$$\mathrm{P}(\{\theta\}, \{\phi\}, data | S_1 .. S_N) \propto \prod_{i=1}^{N} \mathrm{P}(\{\theta\}, \{\phi\}, data \mid S_i) \quad (11)$$

Taking the log of (11), and averaging over sample size N, we get the expected log posterior:

$$\mathrm{L} = \mathrm{E}[\log \mathrm{P}(\{\theta\}, \{\phi\}, \mathrm{data} | S_1 .. S_N)] \approx \frac{1}{\mathrm{NT}} \left(\sum_{\mathrm{i}=1}^{\mathrm{N}} \log \mathrm{P}(\{\theta\}, \{\phi\}, \mathrm{data} | S_i)\right) \quad (12)$$

Evidence Lowe Bound (ELBO) of the model:

$$ELBO[\boldsymbol{p}(\boldsymbol{S})] \coloneqq \boldsymbol{L} - \mathbb{E}[\boldsymbol{log}\, \boldsymbol{p}(S_1 .. S_N)] \quad (13)$$

Train a model to beat a random generator and introduce orthogonality constraint. Model learns from counter examples i.e. random sequences (uniform model).

$$H_T = \kappa[(H_g - H_r) + (H_a - H_r) - H_{cnst}] \quad (14)$$

Constrained expected log-likelihood

$$\ell = \frac{-1}{NT} \left(\sum_{\boldsymbol{i}=\boldsymbol{1}}^{\boldsymbol{N}} H_T^i\right) \quad (15)$$

L2 regularize to avoid overfitting on a large model.

$$L_2 = \lambda \left\| \sum_{j=1}^{q} \sum_{k=1}^{21} \theta_{jk} \right\|^2 + \lambda \left\| \sum_{t=1}^{q} \sum_{j=1}^{q} \sum_{k_2=1}^{21} \sum_{k_1=1}^{21} \varphi_{tjk_1k_2} \right\|^2 \quad (16)$$

Constrained and penalized expected log likelihood and model objective function:

$$\mathcal{L} = \ell + L_2 \quad (17)$$

$$argmin\,(-ELBO) = -\mathcal{L} + \mathbb{E}[\boldsymbol{log}\, \boldsymbol{p}(S_1 .. S_N)] \quad (18)$$

Compare current and previous samples:

$$\Delta(-ELBO) = -ELBO^{i+1} - (-ELBO^{i}) \tag{19}$$

Metropolis-Hastings update allows MCMC sampling:

$$P\left(s^{i} \rightarrow s^{i+1}\right) = \min\left(1,\ e^{-\Delta(-ELBO)}/_{T}\right) \tag{20}$$

Hamiltonian function for HMC, w/ q = {θ, ϕ}:

$$H(p, q) = U(q) + K(p) = -\mathcal{L}(\theta, \phi) + \frac{1}{2} p^{T} M^{-1} p \tag{21}$$

$$p \sim N(0,\ 1) \tag{22}$$

Hamilton's equations of motion:

$$\frac{dq_i}{dt} = \frac{\partial H}{\partial p_i} = [M^{-1} p]_i \tag{23}$$

$$\frac{dp_i}{dt} = -\frac{\partial H}{\partial q_i} = -\frac{\partial U}{\partial q_i} \tag{24}$$

Optimizing the model's stochastic parameters by leapfrog integration of (23) and (24)

$$p_i\left(t + {}^{\varepsilon}/_{2}\right) = p_i(t) - \left({}^{\varepsilon}/_{2}\right) \frac{\partial U}{\partial q_i}(q(t)) \tag{25}$$

$$q_i(t + \varepsilon) = q_i(t) - (\varepsilon) \frac{p_i\left(t + {}^{\varepsilon}/_{2}\right)}{m_i} \tag{26}$$

$$p_i(t + \varepsilon) = p_i\left(t + {}^{\varepsilon}/_{2}\right) - \left({}^{\varepsilon}/_{2}\right) \frac{\partial U}{\partial q_i}(q(t + \varepsilon)) \tag{27}$$

P-value thresholding for top hit prioritization:

$$P(\log(p(S \mid data, \{\theta\},\ \{\varphi\})) \leq X) < threshold \tag{28}$$

**<u>Model's Hyper parameters:</u>**

T: "Temperature"

λ: L2 regularization strength

***κ***: Hamiltonian hyperparameter (balances λ)

Δt: Integration step size (resolution)

L: number of integration steps

N: Training sample per trajectory

M: Number of optimization trajectories

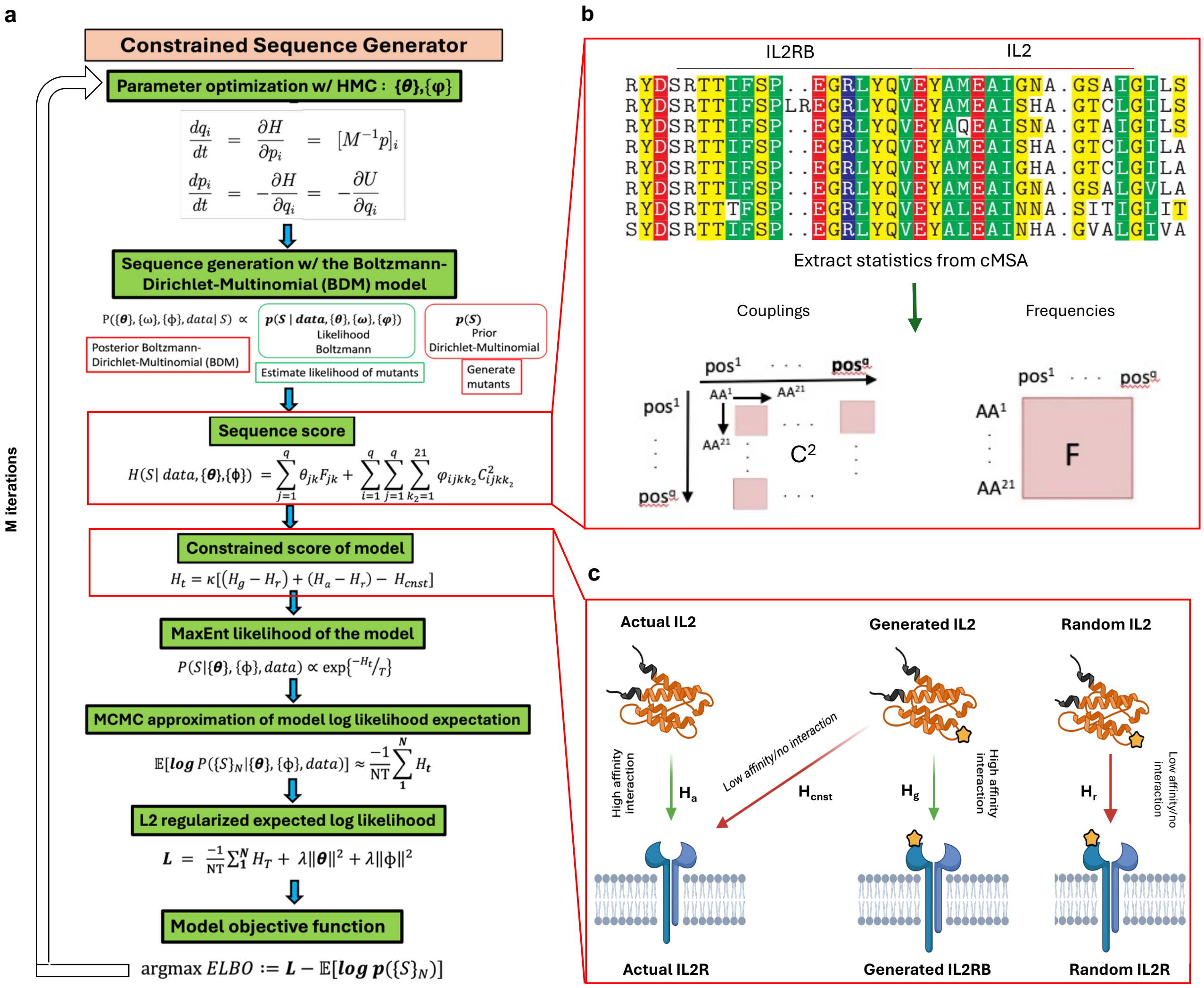


**Supplementary Figure 1. The Constrained Sequence Generator (CSG), a physics-informed machine learning algorithm enabling constrained protein design. a,** Core steps of the training of the CSG algorithm as detailed in Supplementary. **b,** Schematic representation of a cMSA, and the statistics extracted from it, $C^2$, the couplings tensor and, F, the frequency table. **c,** Design concept applied to sequence scoring, different "H" scores reflect the estimated performance of a batch of generated sequence and provide an indication on sample and model quality compared with a random control and reference cMSA (Actual).
